# An open-access neuroimaging-compatible virtual town task for assessing human spatial navigation

**DOI:** 10.64898/2026.11.04.763058

**Authors:** Conor Thornberry, Steven Lieu, James K. Bradshaw Bernacchi, Declan M. Quinn, Lisa Sutton, Rhoda Osagie, Andrew Walsh, Lili Zhang, Tomás Ward, Alejandro López Valdés, Robert Whelan

**Author notes:** Correspondence to: Dr Conor Thornberry, Trinity College Institute of Neuroscience & School of Psychology, Dublin, Ireland.

## Abstract

Spatial navigation is a cognitive skill that draws on multiple processes such as learning, decision-making and memory, which are often updated on a second-by-second timescale. However, many existing virtual paradigms designed to assay spatial navigation cannot target these processes as they unfold or are not designed for integration with concurrent neurophysiological recording. Here, we present and demonstrate our open-access task for neuroimaging: NAturalistic Virtual Assessment of Navigation (NAVAN) combining high-resolution behavioural tracking and integration with electroencephalography. We report data from this task, in which fifty-three participants completed a wayfinding navigation paradigm across twenty-four trials. Navigational efficiency improved significantly following repeated exposure to the environment, although efficiency declined across trials within each block. Self-reported spatial ability (assessed by the Santa Barbara Sense of Direction) predicted performance during early environmental exposure, suggesting concurrent validity. We demonstrated the feasibility of recording electroencephalography during the task, with in-game events streamed and aligned to the neural recording. The task provides researchers with access to detailed behavioural measures and can be adapted by modifying features such as trial number and difficulty. We also provide a step-by-step implementation guide to support reuse and customisation across experimental settings.

## Introduction

Spatial navigation is a cognitive skill we use every day. Though we move through our environments with ease, the underlying neurocognitive processes are complex. We must learn new places, remember old ones, plan routes, orientate ourselves, and correct errors. Spatial navigational ability varies considerably across individuals (Newcombe et al., 2023; Yavuz & Spiers, 2026). The Santa Barbara Sense of Direction scale (Hegarty et al., 2002) is a widely used self-report measure of navigational ability, capturing an individual’s perceived ability to acquire spatial knowledge and estimate distances and directions (Hegarty et al., 2006; Taillade et al., 2016; Ventura et al., 2013). Higher scores have been associated with more efficient navigation (Ventura et al., 2013), suggesting that the scale captures meaningful individual differences in spatial ability. Studying these differences during real-world navigation is challenging because environmental conditions and experiences are difficult to control, making virtual paradigms increasingly valuable for spatial cognition research(Diersch & Wolbers, 2019). Age is one important source of individual variability in navigation. Ageing is associated with changes in spatial navigation and its underlying neurobiology, including a reduced ability to form and use cognitive maps (Coughlan et al., 2018; Iaria et al., 2009; Lester et al., 2017). Furthermore, gender differences have been a prominent research area of individual variability in spatial navigation ability. Although many studies have reported a male advantage on spatial navigation tasks (Astur et al., 1998; Moffat et al., 1998; Nazareth et al., 2019; Padilla et al., 2017), the magnitude of this difference depends on the task and appears to diminish in more naturalistic or real-world navigation settings (Gagnon et al., 2018).

Large-scale tools such as Sea Hero Quest have utilised a gamified approach to assessing spatial navigation. Participants play the game on a mobile device and navigate a boat through virtual environments while completing wayfinding and spatial-memory challenges (Coutrot et al., 2018). Globally, the game has been played by more than four million people worldwide allowing researchers to identify individual differences in navigation abilities linked to age, sex, culture and Alzheimer’s disease risk (Coughlan et al., 2018). Laboratory tasks such as the Virtual Water Maze (VWM) provide translatable, hippocampal sensitive measures of spatial navigation (Commins et al., 2020; Thornberry et al., 2021). Tasks such as the Dual Solution Paradigm (DSP) assays navigation strategy by allowing participants to choose between a learned route and a novel shortcut (Marchette et al., 2011). Despite their utility, these paradigms capture only some aspects of everyday navigation, such as junctions, dead ends, and shortcuts, and many lack compatibility with neuroimaging, which is highly desirable for clinical applications (Whelan et al., 2022). To address these limitations, we developed an openly accessible paradigm, known as the NAturalistic Virtual Assessment of Navigation (NAVAN), which combines naturalistic wayfinding with experimental control and neuroimaging compatibility. Unlike the VWM and DSP, which measure memory for a single hidden goal or route in an open arena, NAVAN requires wayfinding between multiple named landmarks through a structured layout. Participants are not trained along a prescribed route, and can explore the environment freely, select alternative routes, and discover shortcuts without following a previously learned trajectory (Boone et al., 2019).

It is important virtual navigation tasks are compatible with neuroimaging and can record precisely synchronised behavioural data across the full course of navigation. This enables researchers to relate neural activity to specific navigational processes, including planning, movement, shortcut selection, learning and memory (Rolls, 2025; Spiers & Barry, 2015; Stoianov et al., 2018). One prominent account of navigation is cognitive mapping theory (O’Keefe & Nadel, 1978), which suggests we hold internal spatial representations of our environment to guide movement decisions (Tolman, 1948). The hippocampus plays a central role in the formation and use of these representations, as well as in encoding location, distance, direction and spatial relationships (Schiller et al., 2015) (Eichenbaum, 2017). Therefore, accessible and accurate methods that can integrate with neuroimaging are required to systematically model and test these current theoretical frameworks. Navigation also depends on distributed cortical systems (Beest et al., 2026). The frontal cortex is thought to be involved in spatial attention, motor planning and working memory (Delaux et al., 2021). Theta oscillations (4-8 Hz) are brain waves thought to coordinate these networks and have been shown to increase in power during virtual navigation (Chrastil et al., 2022; Thornberry & Commins, 2024). A robust finding is that frontal midline theta increases are linked to the onset of active spatial navigation in virtual tasks (Chrastil et al., 2022; Du et al., 2023). In addition, theta power has been shown to increase in the human hippocampus during virtual movement (Ekstrom et al., 2005). Hence, brain oscillations within the theta range, which when recorded from the hippocampus can be as broad as 1-10Hz (Jacobs, 2014), are a vital neural component of human navigation (Buzsáki & Moser, 2013; Herweg et al., 2020). Accordingly, methods that align continuous behavioural measures and task events with neural recordings are essential for testing mechanistic accounts of human spatial navigation.

Therefore, to demonstrate the feasibility and capabilities of our virtual town for navigation experiments (NAVAN), we outline three primary hypotheses based on the existing literature that will support the use of this task. First, we hypothesise that individual differences in self-reported spatial ability, as measured by the SBSOD scale, will predict task performance, such that higher scores are associated with more efficient navigation. Second, we predict improved navigational performance following repeated exposure to the environment, such that performance improves across blocks of trials. Third, we assess the feasibility of recording and analysing neural oscillatory activity during the task in a small EEG subsample. As an exploratory prediction, we expect differences in theta-band power (5-7 Hz) between the planning and active navigation phases, consistent with the established role of frontal theta in active spatial cognition and navigation (Du et al., 2023; Thornberry & Commins, 2024).

## Methods

### Participants

A total of 59 participants were recruited across two experiments. Experiment 1 investigated behavioural performance only, whilst Experiment 2 examined the feasibility of using the gamified task with EEG (n=6). Experiment 1 included 53 participant (33 females, 20 males) with mean age of 21.71 years (SEM +/- 0.101). In our EEG experiment, our sample contained usable data of n = 6 (6 female; age range 20-30 years).

### Santa Barbara Sense of Direction Scale (SBSOD)

The SBSOD is a standardised, paper-based self-report scale of spatial ability (Hegarty et al., 2002). The assessment consists of 15 statements describing spatial behaviours, preferences, and perceived abilities in real-world navigation such as “I am very good at giving directions”. Participants rate the extent to which each statement applies to them using a 7-point scale ranging from 1 (strongly agree) to 7 (strongly disagree). After reverse-scoring the appropriate items, higher mean scores indicate a better self-reported sense of direction, and lower scores indicate a poorer sense of direction.

### Virtual Navigation Task

The experimental paradigm consisted of a spatial navigation task in which participants navigated between landmarks in a 3D virtual town environment (Figure 1A) under a fixed time constraint (30 seconds). The virtual environment was developed in Unity and displayed on a 26-inch monitor at 1060 x 1080 resolution and 100 Hz refresh rate. Navigation was performed using a standard gamepad controller with the left analogue stick controlling movement direction and the right analogue stick controlling viewing direction. The virtual town measured 200 × 200 virtual metres. The environment comprised 10 × 10 virtual-metre (Vm) tiles (Figure 1B), with 81 navigable tiles providing 8,100 Vm² of navigable space (Figure 1C). There is a total of nine “destinations” (landmarks) available to visit by the player, each with a unique name and sign (see Figure 1C). The destinations are marked by 3D flat square tiles on the floor. They are (1.5 × 1.5 × 0.05 Vm) metres in size. All task documentation, datasets, an easy-to-use executable version of the task itself as well as the core Unity code are available via Zenodo [Link Following Acceptance]. Further details are provided in the section “Using the Virtual Town for Navigation Experiments”.

**Figure 1:**
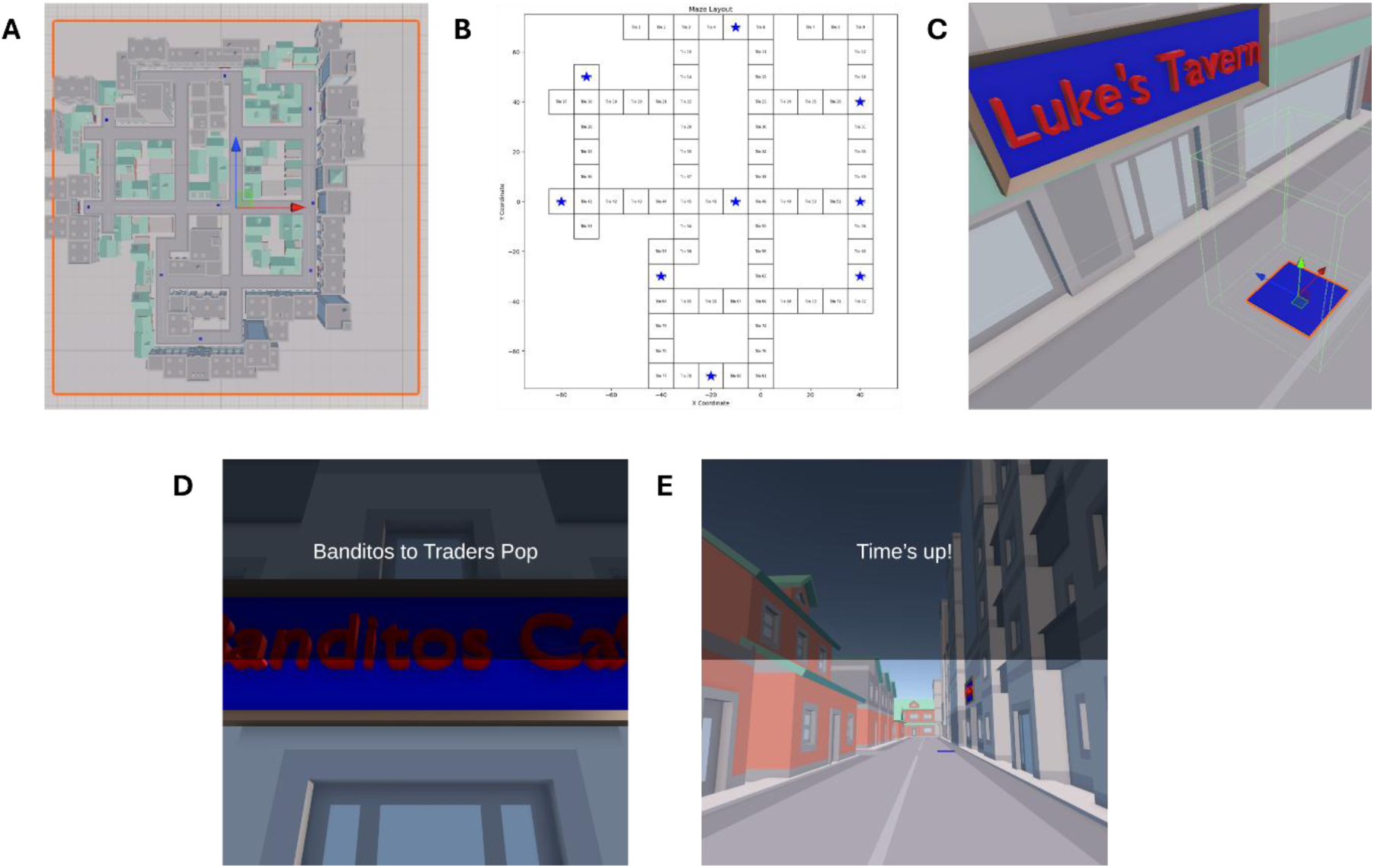
Task Design & Layout. (A) Overhead view of the wayfinding environment. (B) Tile layout with landmark locations. (C) Target square design with landmark/target signage showing destination. (D) Screenshot from the participant’s perspective of the planning prompt before a trial begins. (E) Screenshot from the participant’s perspective of the “Time’s up!” notification of a failed trial, which restricts movement and the participant is relocated to the correct location.

### Experimental Paradigm

Prior to the navigation task, participants completed a 5-minute practice period during which they familiarised themselves with the virtual environment and gamepad controls. Participants were allowed to explore the town freely with the landmarks visible but were not instructed to memorise routes or landmark locations.

At the start of the navigation task, participants were placed at an assigned starting landmark (Table 1). A grey message box presented at the centre of the screen instructed participants to navigate to a named destination (e.g. “Gemma to Banditos Café”). Participants were informed that destinations should be reached as quickly and accurately as possible, and that no map of the environment would be provided. During this initial stage, participant movement was disabled for 10 s, marking the planning phase, during which participants were able to plan their route before movement onset. The destination instruction remained visible throughout this period (see Figure 1D). Once the 10-s interval elapsed, participants had a 30-s time limit to navigate to the named target, which began once the text box disappeared. Arrival at the destination was registered when the participant moved onto a blue square positioned at the target, which stopped the timer and triggered the onset of the next trial. Participants were then given a subsequent destination prompt. If a participant was unsuccessful in locating the destination within the 30-s time limit, they were automatically relocated to the correct destination for 3 s prior to the onset of the next trial, providing brief passive exposure to the target location. No performance feedback was given on a trial-by-trial basis, but participants were alerted that their trial was over by a “Time’s Up!” notification (Figure 1E).

**Table 1:** Euclidean inter-landmark distances (nearest Vm) in the virtual city environment.

| **Landmark** | **1** | **2** | **3** | **4** | **5** | **6** | **7** | **8** | **9** |
|---|---|---|---|---|---|---|---|---|---|
| **1. Lukes Tavern** |  |  |  |  |  |  |  |  |  |
| **2. QBit Bank** | 50 |  |  |  |  |  |  |  |  |
| **3. RINO Corp** | 84 | 51 |  |  |  |  |  |  |  |
| **4. Banditos** | 115 | 87 | 36 |  |  |  |  |  |  |
| **5. Joeys** | 142 | 138 | 96 | 71 |  |  |  |  |  |
| **6. Cafe Zondag** | 112 | 121 | 92 | 82 | 41 |  |  |  |  |
| **7. Traders Pop** | 103 | 133 | 123 | 126 | 94 | 54 |  |  |  |
| **8. Gemma** | 69 | 112 | 121 | 138 | 129 | 89 | 48 |  |  |
| **9. Lumiere** | 70 | 73 | 52 | 62 | 73 | 49 | 71 | 77 |  |

*NOTE: Values are Euclidean distances between landmark pairs.*

The task consisted of two levels, each containing twelve trials, presented within the same virtual environment such that spatial knowledge accumulated continuously across levels. Trial order within each level was held constant across participants to yield a comparable learning rate within the sample. Both levels were designed to balance inter-landmark path lengths across trials (Table 1). The optimal travel path length between each pair of landmarks was calculated *a priori* from the layout of the navigable environment (Figure 1B). Level 2 was differentiated from Level 1 by introducing novel starting points for targets navigated to in Level 1. Mean optimal path length was closely matched between Level 1 (M = 121.884 Vm, SD = 16.04 Vm) and Level 2 (M = 121.91 Vm, SD = 17.48 Vm) is maintained.

### EEG Acquisition & Preprocessing

EEG data were acquired using a BioSemi ActiveTwo system (BioSemi B.V., Amsterdam, Netherlands) with 128 Ag/AgCl electrodes positioned in the radial ABC system layout, recorded at a sample rate of 512 Hz. Continuously recorded EEG data were synchronised to behavioural events using Lab Streaming Layer (Kothe et al., 2025) producing a single .XDF file. Implementation details are provided in the section “Using the Virtual Town for Navigation Experiments”.

EEG data were then analysed offline in MATLAB R2025 using scripts within the Brainstorm package (Tadel et al., 2011). A 1-Hz high-pass filter and a 40-Hz low-pass filter were applied. Data were visually inspected for bad segments and bad electrodes, which were then removed. Independent component analysis (ICA), using the EEGLAB infomax algorithm, was performed on the data. Components were then manually inspected to identify artefacts, which were subsequently removed. Data were epoched from −1000 ms to +4000 ms after relevant event markers (Planning and Start). Epochs with voltage steps above 100 μV or peak-to-peak signal deflections exceeding 200 μV were rejected. EEG data were re-referenced to an average of 128 scalp electrodes. EEG data were epoched around two events: planning (target instruction onset, movement disabled) and active navigation (movement onset), across −1000 ms to +4000 ms from event onset. Welch Power Spectrum Density (PSD) computed for 1-40 Hz over 0 ms to 3000 ms (selected as a period of interest to capture initial virtual movement onset and active navigation) with a 1-sindow and a 50% overlap. Mean theta power was then extracted at 5-7 Hz based on the default definition in Brainstorm (Tadel et al., 2011); this range also avoided overlap with the predefined delta (2–4 Hz) and alpha (8–12 Hz) bands.

### Outcome Measures, Statistical Analysis and Ethics

Behavioural outcomes included navigation duration, path length, path efficiency, standstill count, and success rate. *Navigation duration* was measured in seconds from movement onset to destination arrival. *Path length* was the total distance travelled in virtual metres. Path *efficiency* was calculated as the proportion of optimal to actual path length (in virtual metres), where a score of 1 indicates a perfectly efficient trajectory. *Standstill count* was the number of instances where x and y coordinates remained unchanged for a minimum of four consecutive samples (i.e., ≥1 s). *Success rate* was the proportion of trials completed within the 30-s limit.

Paired-samples *t*-tests examined differences between levels. A Linear Mixed Model was used to examine Level and Trial effects on path efficiency as well as the interaction between Level and Trial using JASP version 0.96 (Love et al., 2019). Fixed effects were Level (within-participant factor with two levels) and Trial (entered as a mean-centred continuous linear predictor), together with their interaction. Participant gender was included in the model as a covariate considering well-documented gender differences in navigation. Path efficiency was selected as the primary trial-level outcome because it accounts for variation in the optimal distance between starting points and goals. Pearson correlations were used to examine bivariate associations with SBSOD scores, followed by a hierarchical regression to examine SBSOD relationships. EEG analyses used paired permutation *t*-tests within Brainstorm. The study and task were approved by the School of Psychology Research Ethics Committee, Trinity College Dublin (#3832).

## Behavioural Results

### Predicting task performance from SBSOD

To examine whether self-reported spatial ability predicted navigation task performance beyond demographic variables, a hierarchical regression was conducted with path efficiency from Level 1 as the outcome variable, as earlier navigation likely indexes environmental encoding. Initially, correlations were used to explore the relationship between all available variables and SBSOD, and to check for multicollinearity. The results of these are available in Supplementary Materials. Gender and age were entered in Model 1, which did not explain a significant proportion of variance in path efficiency (R² = 0.035, *F*_(2, 50)_ = 0.916, *p* = 0.407). Neither gender (β = 0.029, *t* = 1.182, *p* = 0.243) nor age (β = −0.013, *t* = −0.808, *p* = 0.423) were significant predictors. SBSOD scores were entered in Model 2, producing a significant increase in explained variance (R² change = 0.179, *p* = 0.002). The full model accounted for 21.4% of variance in Level 1 path efficiency (R² = 0.214, adjusted R² = 0.166, *F*_(3, 49)_ = 4.448, *p* = 0.008). SBSOD was the only significant predictor (β = 0.461, *t* = 3.338, *p* = 0.002), indicating that participants who reported higher spatial ability navigated more efficiently during initial spatial navigation. A second hierarchical regression with Level 2 path efficiency examined if SBSOD predicted relearning. Gender and age were entered into Model 1, which did not significantly explain the variance in path efficiency (R² = 0.058, *F*_(2, 50)_ = 1.552, *p* = 0.222). Neither gender (b = 0.049, *t* = 1.756, *p* = 0.085) nor age (β = −0.002, *t* = −0.091, *p* = 0.928) were significant predictors. SBSOD scores were entered in Model 2, producing an increase in explained variance that did not reach statistical significance (R² change = 0.035, *p* = 0.174). Thus, SBSOD scores predicted path efficiency during initial exposure but not following re-exposure to the environment.

### Spatial Navigation Improvements Across Levels

Paired-samples t-tests compared behavioural performance between Level 1 (L1) and Level 2 (L2) across the 53 participants. Descriptive statistics and test results are summarised in Table 2. Performance improved across all measures from Level 1 to Level 2 (all *p* < 0.001). Trial duration decreased by an average of 2.36 s (Cohen’s *d* = 0.71), and path length decreased by 15.28 virtual metres (Cohen’s *d* = 0.69). Standstill count also decreased (Cohen’s d = 0.36). Path efficiency increased from 0.63 to 0.70 (*t*(52) = −8.08, *p* < 0.001, Cohen’s d = −0.75), and success rate increased from 73% to 85% (*t*(52) = −5.08, p < 0.001, Cohen’s d = −0.48) (Figure 3A-D).

**Figure 3:**
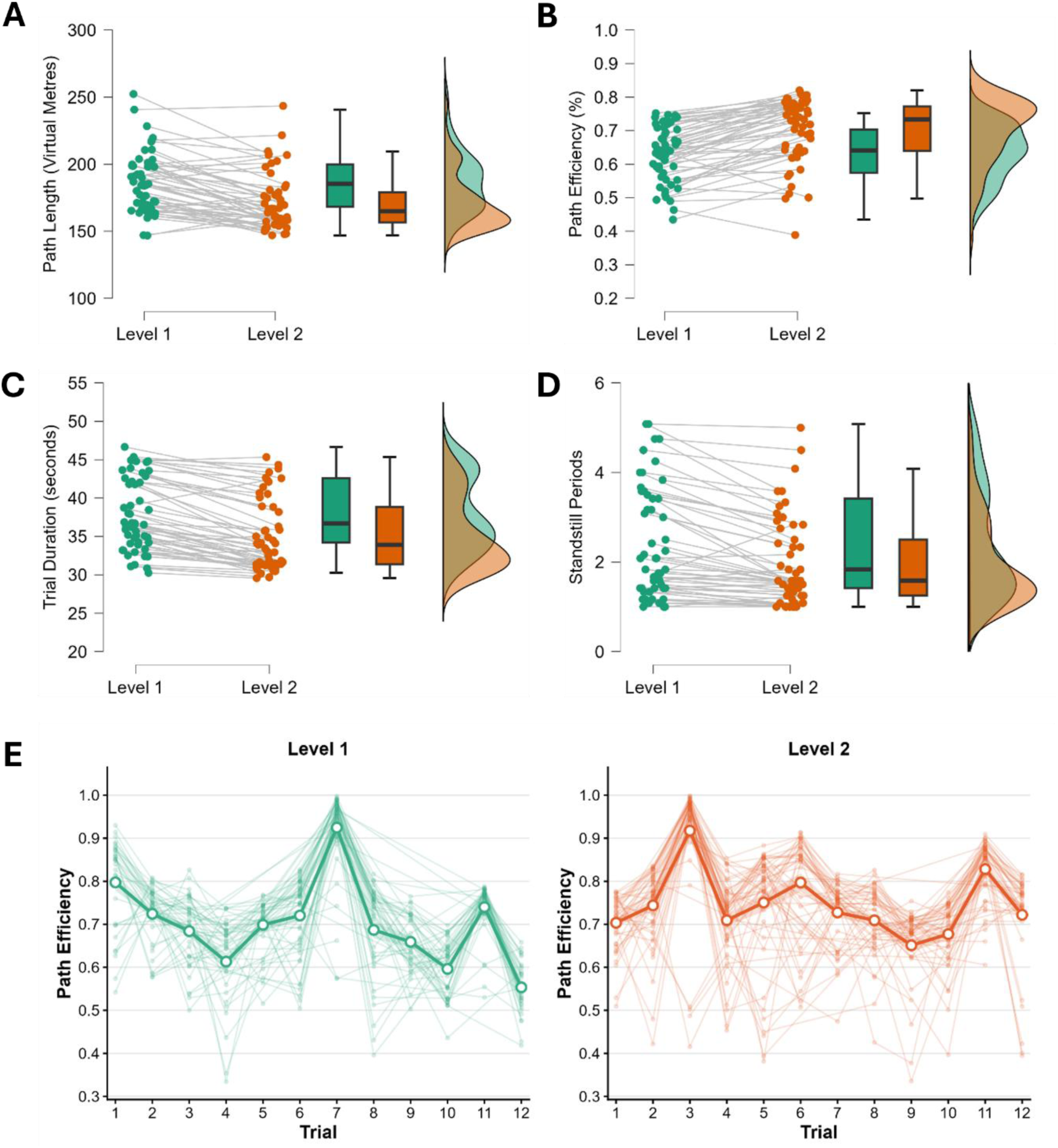
Behavioural performance across levels and trials. Paired individual data points with box plots and marginal density distributions for each dependent variable: path length in virtual metres (A), path efficiency as a percentage (B), trial duration in seconds (C), and standstill periods (D). Green and orange data points represent Level 1 and Level 2 respectively, with grey lines connecting individual participants across levels. (E) Trial-by-trial path efficiency across the 12 trials of Level 1 (green) and Level 2 (orange). Individual participant trajectories are shown as faint lines, with the group mean trajectory indicated by the bold line.

**Table 2:** Descriptive statistics and paired-samples t-test results of behavioural measures of all trials (successful and unsuccessful). L1 and L2 are the two game levels (Level 1 and Level 2), with M (mean) and SD (standard deviation) for each variable of interest. Cohen’s d and the 95% Confidence Interval (CI) are provided for each statistical test.

|  | L1 *M* (SD) | L2 *M* (SD) | *t* | *p* | *d* | 95% CI |
|---|---|---|---|---|---|---|
| Duration (s) | 22.07 (3.39) | 19.71 (3.18) | 9.01 | <0.001 | 0.71 | [0.41, 1.01] |
| Path Length (Vm) | 186 (22.37) | 171 (21.31) | 6.84 | <0.001 | 0.69 | [0.40, 1.00] |
| Path Efficiency | 0.63 (0.08) | 0.70 (0.10) | -8.08 | <0.001 | -0.75 | [-1.06, -0.44] |
| Standstill Count | 2.40 (1.24) | 1.96 (0.99) | 5.7 | <0.001 | 0.36 | [0.08, 0.63] |
| Success Rate | 0.73 (0.28) | 0.85 (0.23) | -5.08 | <0.001 | -0.48 | [-0.76, -0.19] |

Trial-level analyses were then restricted to successful trials; resulting in and exclusion rate of 20.9%. Random effects included participant ID to account for interindividual differences; the random-effects structure produced a singular fit, so the model was reduced to random intercepts. The model was estimated using restricted maximum likelihood, and degrees of freedom were calculated using the Satterthwaite method. Type III sums of squares were used. The model was fitted to 1,006 observations from 53 participants. Path efficiency was significantly higher in Level 2 than Level 1, *F*_(1, 964.22)_ *=* 31.02*, p <* 0.001 (See Figure 3B & 3E). For Level, estimated marginal means were 0.70, 95% CI [0.69, 0.72] for Level 1 and 0.75, 95% CI [0.73, 0.76] for Level 2, a difference of 0.05, 95% CI [0.03, 0.06], *t*(964.22) = 5.57, *p* < 0.001. Participants therefore navigated closer to the optimal route on their second exposure to the environment. Path efficiency declined across successive trials within each level, *F*_(1, 955.61)_ = 35.78, *p* < .001, *b* = −0.007, SE = 0.001, *t*(955.61) = −5.98, a reduction of approximately 0.7 percentage points per trial. This decline differed by Level (Trial X Level interaction), *F*_(1, 956.70)_ = 3.997, *p* = 0.046, *b* = −0.002, SE = 0.001, *t*(956.70) = −1.99. Gender did not significantly predict path efficiency, *F*_(1, 44.08)_ = 0.002, *p* = 0.965.

### Gender Differences

Independent samples *t*-tests examined whether male and female participants differed on task performance across both levels. Male participants had significantly shorter trial durations than female participants in both Level 1 (Male: *M* = 20.73s, *SD* = 3.39s; Female: *M* = 22.82s, *SD* = 3.2s; *t*(51) = 2.231, *p* = 0.03, *d* = 0.639) and Level 2 (Male: *M* = 18.31s, *SD* = 2.60; Female: *M* = 20.49s, *SD* = 3.23s; *t*(51) = 2.52, *p* = 0.015, *d* = 0.722). Male participants also paused significantly less frequently than females in Level 1 (Male: *M* = 1.605, *SD* = 0.916; Female: *M* = 2.846, *SD* = 1.173; *t*(51) = 3.975, *p* < 0.001, *d* = 1.139) and Level 2 (Male: *M* = 1.373, *SD* = 0.558; Female: *M* = 2.292, *SD* = 1.025; *t*(51) = 3.610, *p* < 0.001, *d* = 1.034). There were no significant differences in path efficiency in either level (Level 1: *t*(51) = −1.089, *p* = 0.281; Level 2: *t*(51) = −1.777, *p* = 0.082), indicating that while male participants navigated with greater speed and less hesitation, they did not demonstrate greater efficiency compared to female participants. See Figure 4 for a summary of these results. Finally, male participants reported significantly higher SBSOD scores (M = 4.635, SD = 1.09) than female participants (M = 3.776, SD = 1.015; *t*(51) = −2.88, *p* = 0.006, *d* = −0.824).

**Figure 4:**
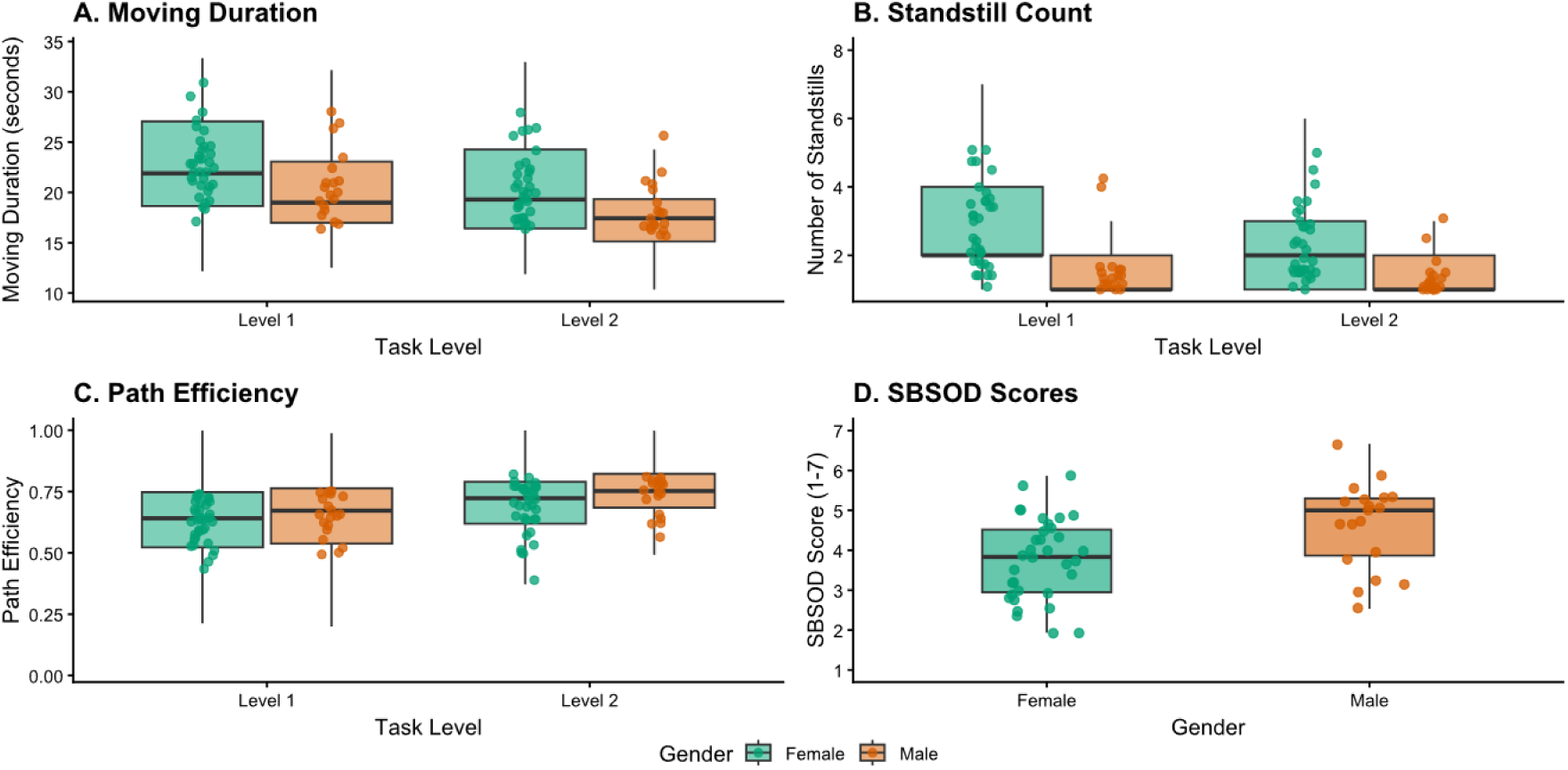
Boxplots showing mean (A) Trial Duration, (B) Standstill Count and (C) Path Efficiency for males and females during level 1 and level 2. Individual means are marked by jittered points. (D) A boxplot showing average Santa Barbara Sense of Direction (SBSOD) with mean scores jittered for males and females.

## EEG Feasibility

### Theta differences between planning and navigation

Given the small EEG sample, analyses were intended primarily to demonstrate feasibility rather than provide confirmatory tests of theta effects. We first conducted exploratory comparisons of theta power (5–7 Hz) between Level 1 and Level 2 separately for the planning and active-navigation phases. Uncorrected paired permutation t-tests showed no significant differences in theta power were observed between levels during either the planning or active navigation phases (Figure 5A-B), nor between planning and active navigation when analysed separately by level (all *p* > 0.05, see Figure 5C-D). For the principal feasibility comparison, PSD values were then averaged across trials and levels for each participant,, and all statistical tests were conducted on these participant-level averages (n = 6). With six participants, a paired sign-flip permutation test has 64 possible permutations, so the smallest attainable two-tailed p-value is approximately 0.031. As this exceeds any FDR-corrected threshold across 128 electrodes, results are reported uncorrected. Mean theta (5-7 Hz) power during planning and active navigation is shown in Figures 5E and 5F. Theta power was greater during active navigation than during planning at 5 electrodes (uncorrected *p* ≈ 0.031, Figure 5G). These formed three groups of electrodes: a central-frontal group (approx. AFz and C6), and a posterior group (BioSemi A24, A27, A29). These results demonstrate that task-locked EEG can be recorded and analysed during the task, rather than providing evidence of any theta power difference. This is evidence that the task is capable of recording theta oscillations during active navigation using stimulus-locked analyses of naturalistic navigation.

**Figure 5:**
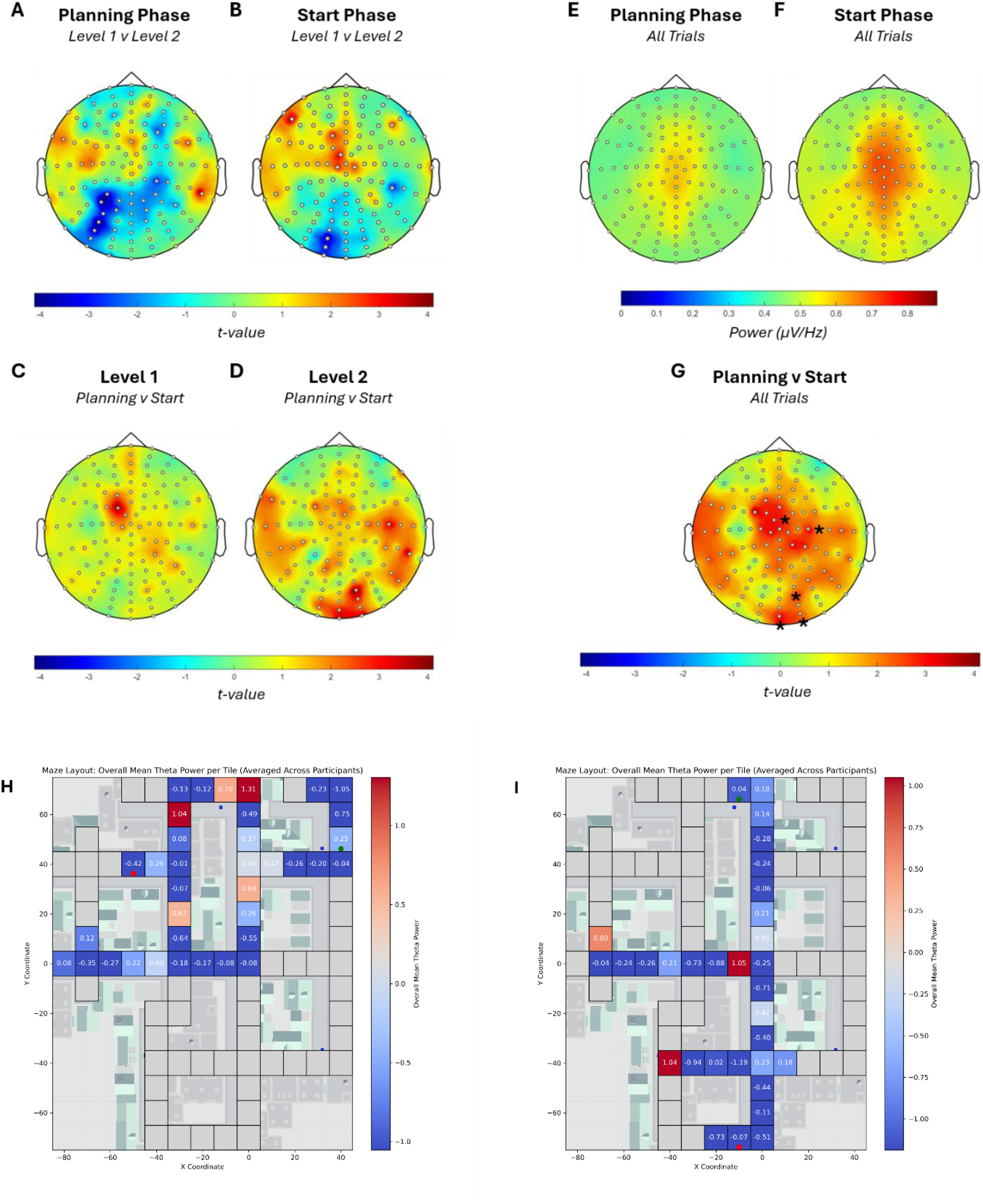
EEG Results. Group-level permutation *t*-test contrasting theta power (5-7 Hz) between levels during the planning (A) and active navigation (B). Between-phase comparisons within Level 1 (C) and Level 2 (D). Grand-average theta power (µV²/Hz) collapsed across all trials and levels, for the planning (E) and active navigation (F). (G) Permutation *t*-test results for planning vs. active navigation across all trials and levels; asterisks denote statistically significant differences. Colour scales indicate *t*-statistics (A-D, G) or power magnitude in µV²/Hz (E-F). Path travelled and theta power overlays across participants for Trial 1 (H) and Trial 12 (I) of Level 1, demonstrating step-by-step neural dynamics during navigation.

### Additional Analysis Capabilities

The 4 Hz sampling rate allows step-by-step behavioural dynamics (recorded every 250 ms) to be mapped spatially across the grid environment, synchronised with EEG recordings via Lab Streaming Layer (Kothe et al., 2025). Example group-averaged route-theta power overlays for Trial 1 and Trial 12 of Level 1 Figure 5H–I shows illustrative group-averaged overlays of route position and theta power for Trial 1 and Trial 12 of Level 1. These visualisations are provided for demonstration purposes and do not constitute inferential analyses. Code and reusable building blocks for generating such plots repository are available on the Zenodo repository.

## Using the Virtual Town for Navigation Experiments

### Task Overview

From the main menu, the experimenter can open the Virtual Town task and configure the settings for a particular experimental paradigm. After configuration, selecting Start places the participant in the town and presents a target destination on screen. By default, the participant has 30 seconds to reach each target, although this duration can be changed. Successful arrival triggers the next target; if the target is not reached within the allotted time, the participant is relocated to the target before the next trial begins. This sequence continues until all configured destinations have been presented. At task completion, a summary screen is displayed, after which the task can be repeated or exited.

### Menu Navigation

When the task is started via the executable file (.exe), the main menu screen (Figure 6; Left) will be opened where the experimenter is presented with several options. The *Player Scene* and *Player vs Agent Scene* will load the navigation game mode where the player must find a cube in the maze. This mode is experimental and is intended for collecting data to train AI agents in the maze as part of a separate project. However, researchers are more than welcome to contribute to development and testing in this area. *Wayfinding Test (WFT) Mode* loads the Virtual Town task used in this study. Replay Scene allows previously saved files to be loaded and replayed, and Exit closes the application. During gameplay, the pause menu can be opened by pressing the Escape key on a Windows keyboard (see Figure 6; Right). The *Resume* button will close the pause menu and resume gameplay. The *Reset Scene* button will exit the current loaded task and reload the game options menu (discussed below). This has the same effect as selecting *Wayfinding Test Mode* from the main menu. This option is useful if after completing an attempt at one of the game modes, the experimenter wishes to try again with different options or restart the recording. The *Return to Main Menu* will load the main menu. Finally, the *Exit Game* button will close and exit the game.

**Figure 6:**
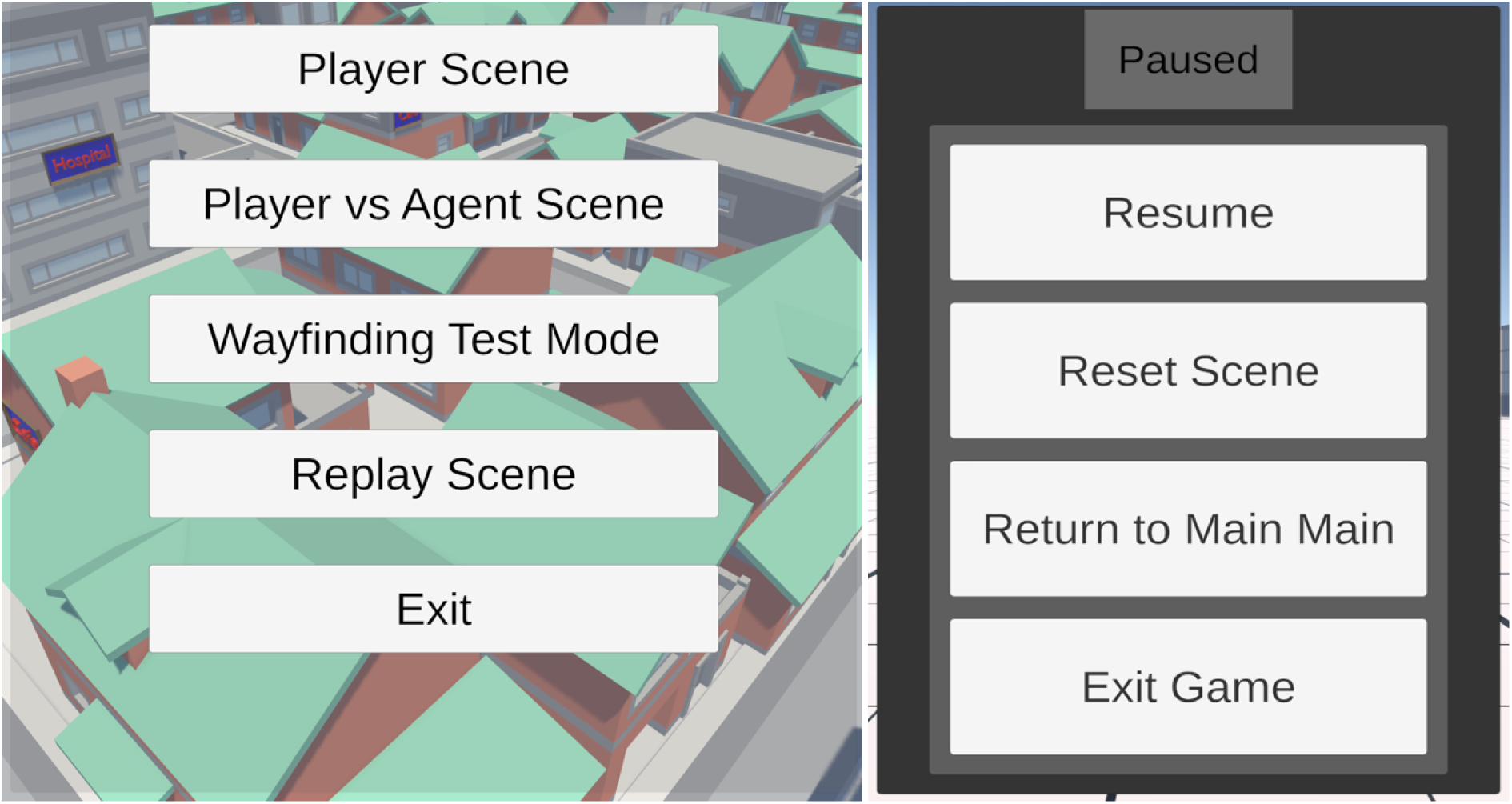
Main Menu and Pause Menu of the Virtual Town Task. The main menu (left) is displayed when the application is launched. Player Scene & Player vs Agent Scene: Loads an experimental game mode where the player searches for a cube in the town; designed for gathering data to train AI agents. Wayfinding Test (WFT) Mode: Loads the primary Virtual Town task used for the experimental spatial navigation testing. Replay Scene: Opens a scene where previously saved task files can be loaded and replayed. Exit: Closes and exits the application. On the right, this menu will appear during the task if the experimenter or the player presses the Esc key, referred to as the game being paused. Resume: Closes the pause menu and resume active gameplay. Reset Scene: Exits the current task and reloads the game options setup menu (see below), allowing you to restart or adjust settings without returning to the title screen (Left). Return to Main Menu: Exits the active session and returns to the main menu screen. Exit Game: Immediately closes and exits the application.

### Player Controls and Accessibility

The player can control the character in the navigation task via a keyboard and mouse plugged into a Windows PC. However, a Windows compatible gamepad (such as a Microsoft Xbox controller) may be connected to the PC and will automatically work with the task. Other joystick or controller types have not been systematically tested and may require external button mapping to reproduce the W, A, S, and D inputs used by the task.

Keyboard:

- Move forward: **W**
- Move backward: **S**
- Move left: **A**
- Move right: **D**
- Rotation: **Mouse movement**

Controller:

- Move forward: Left analogue stick forward
- Move backward: Left analogue stick backward
- Move left: Left analogue stick left
- Move right: Left analogue stick right
- Rotation: Right analogue stick movement

Menu / User Interface (UI) Navigation:

All the UI buttons and options are only accessible with a mouse. Pause menu:

- Keyboard: **Escape**
- Controller: **Start Button** or **Option Button**

### Task Configuration

#### Level Select

Two levels, corresponding to those used in the present study, are included with the WFT. Each level has its own set of destinations for the player to reach. The levels are played on the same map, with only the destinations changing (Figure 7A). *Training Mode:* Enabling this option disables the WFT destinations, allowing the player to navigate through the maze with no time limits or targets. The player can familiarise themselves with the layout of the maze and the chosen controller/gamepad setup using this option (Figure 7B). Player-position data are recorded while training, no events markers are recorded (no destinations to trigger the events). Training mode can be exited through the pause menu by selecting Reset Scene. *Stream Type*: Two Lab Streaming Layer stream types are available for behavioural data: Marker and Signal (Figure 7C). In general, **a marker stream should be used** (as this records X and Y coordinates every 250 ms which can be combined with the EEG data). However, a signal stream is *required* if using the LSL data with OpenViBE; a brain-computer interface (Renard et al., 2010). Note that this compatibility is preliminary and may be developed by researchers should they wish to integrate brain–computer interface’s into the task. The stream choice **does not** affect the conditional events in the task. These events will always be built into your LSL stream when the game is active. Conditional task events are included regardless of the selected stream type. To initialise LSL **before** the task begins, select the button beside LSL Status (Figure 7D). The status changes to green (“Open”) when the selected streams are detected. This allows compatible recording software, including BioSemi acquisition software or OpenViBE, to connect to the LSL streams before task onset. Selecting Start (Figure 7E) begins the task using the configured options; if the streams have not already been initialised, they start automatically at this point.

**Figure 7:**
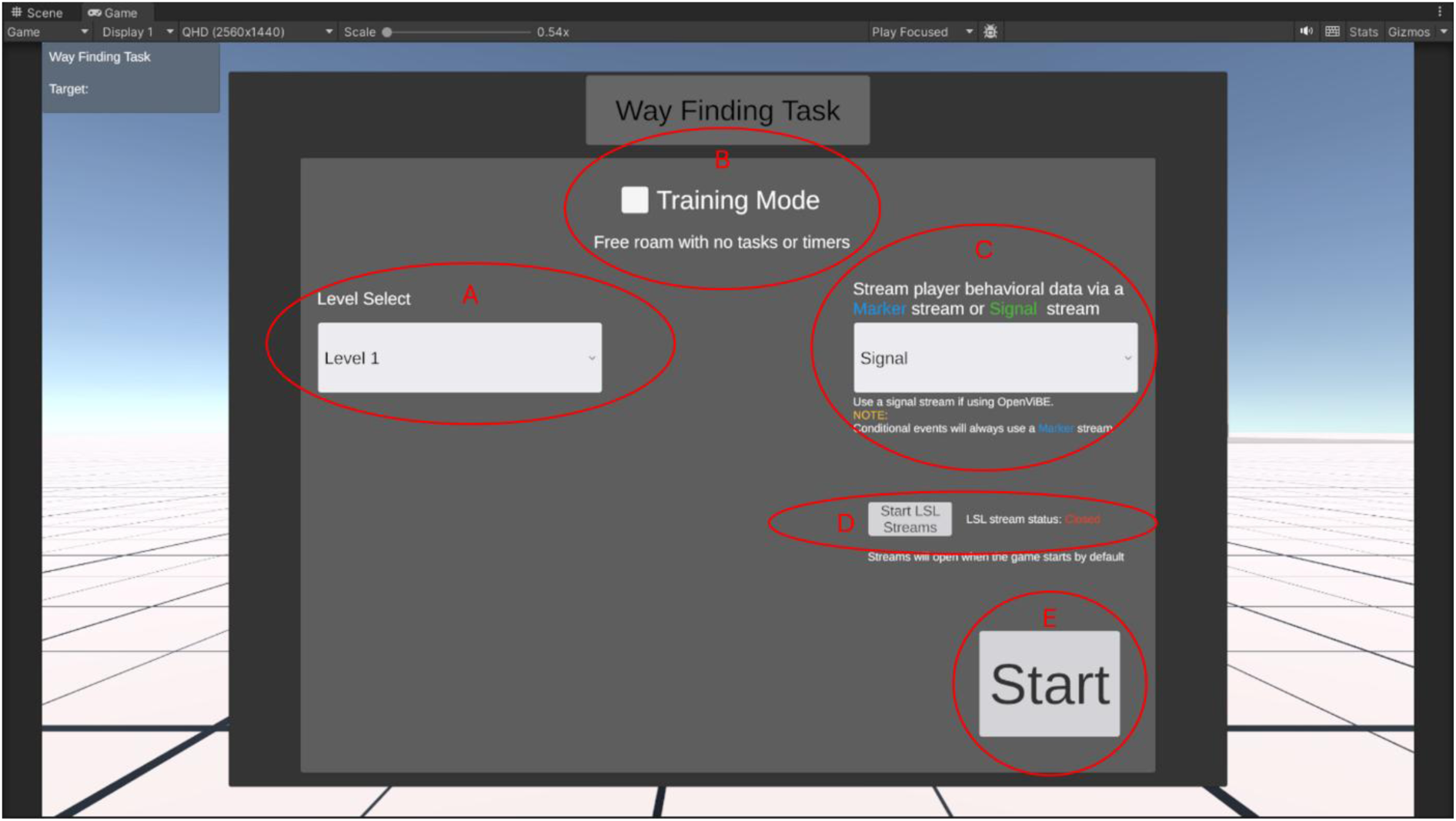
Main task configuration screen for the Way Finding Task. From this screen the experimenter can select the Level (A), turn on or off Training Mode (B), select the type of event streaming they would like to use (C), start the event streaming (D) and finally start the task by pressing the Start button (E).

### Further Details on LSL Data Streams

Player behavioural data and conditional events are streamed via Lab Streaming Layer (LSL). When the player data is recorded and stored in memory, the data is pushed into an LSL stream. This LSL data can then be received and read by any software that supports LSL. There are 3 LSL data streams in the WFT: (1) Player data stream; (2) Conditional events stream and (3) Continuous player data stream.

*The player data streams will appear in Lab Recorder as the following:*

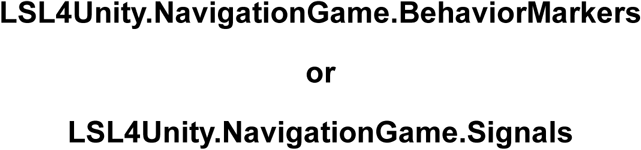

The player data stream includes: (1) The XYZ position of the player in the maze (three values). (2) The XYZ rotation of the player in the maze (three values). (3) Distance to the current target (one float value). (4) Step count (one integer value). (5) Time stamp (only included in marker stream as one string value).. Player data can be streamed in either marker or signal format. The marker stream uses string-formatted values, allowing heterogeneous data types to be transmitted and subsequently parsed. The signal stream uses floating-point values and is intended for use with OpenViBE.

*The conditional events stream will appear in Lab Recorder as the following*:

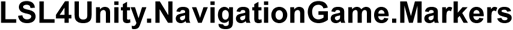

The following events are streamed as integers:

1. **Level Start**: The participant is oriented to face the sign marking the initial landmark (“Luke’s Tavern” in Level 1 or “Gemma” in Level 2). This event marks the start of the level, not the start of each individual trial.
2. **Planning Phase**: The instruction screen appears as a black rectangle at the centre of the display and presents the route instruction specified by the selected level (e.g., “Gemma to Café Zondag”). The instruction remains visible for 10 s while movement controls are locked. This event marks the beginning of the planning phase used in the present experiments.
3. **Active Navigation**: The instruction screen disappears, movement controls are unlocked, and the participant can begin moving. This event marks trial onset and the start of the active-navigation phase. The 30-s time limit begins at this point.
4. **Failed Trial**: This event is sent if the participant does not reach the destination within the time limit. The event is generated when the trial ends, the “Time’s Up!” message and movement is once again restricted.
5. **Successful Trial**: This event is sent when the participant reaches the destination by entering the target area (blue square), at which point movement is restricted and the trial ends.

*The continuous player data stream will appear in Lab Recorder as*:

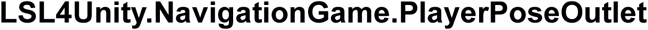

This data stream is similar to the *player data stream* with the exception that the data in this stream is sampled at a much higher rate than the previous streams and conditional event markers listed above. Data are pushed into the stream at the update rate of 50 frames per second and is only streamed as a signal stream for use with OpenViBE. This stream contains only the local XYZ position and local XYZ rotation of the player in the maze.

### Data Management

Player data is recorded in two ways: save files (.json) and LSL data streams (.xdf). As the player plays the game, their data is recorded and written to JSON. When the player finishes the WFT the data is written to a JSON save file. There are 2 types of data recorded: game session data, and player step data. The game session data includes Mode (training vs. non training); Map name; Duration of the task; Number of successful locations reached; Number of unsuccessful locations reached. The player step data is recorded at regular intervals (4 Hz) and includes: Step count (used for the **replay system**, see below); Distance to the current target; Time stamp; The X Y position of the player in the town and the rotation of the player in the town. The current target. An example of the JSON file structure is displayed below in Figure 8. The save files are written to this JSON file when the task is completed or if the task is exited early via the pause menu by doing one of the following: Resetting the scene; Exiting the scene to the main menu or exiting the game. This means that should an error occur during data collection, the data up to the point of that issue will be saved. The save files can be found at a data path on the Windows system being used. The initial AppData folder will have to be made visible by selecting “Unhide” in your properties or in the File Viewer selecting View -> Show -> Hidden Items. The typical path containing the files (Figure 8) is the following:

**Figure 8:**
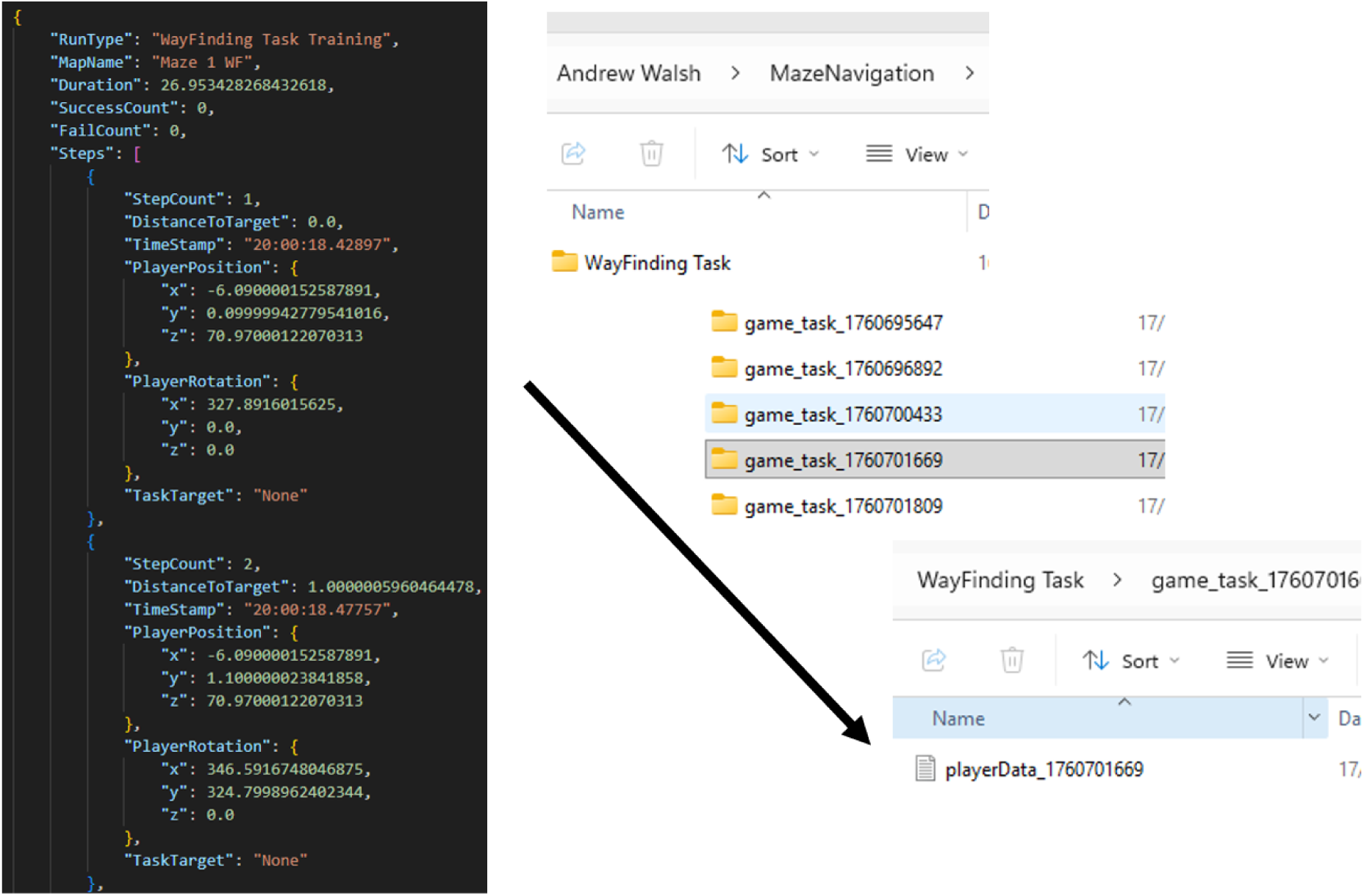
File and Folder Outputs. (Left) Structure of task results in JSON file outputs. (Right) Folder structure in which JSON file outputs are found.

C:\Users\<user>\AppData\LocalLow\User_name\MazeNavigation

### Replaying Saved Files

The JSON save files generated by the player can be loaded and replayed in the “Replay Scene” mode, which can be found on the main menu. Copy the save file path and load the file. This may be useful for reconstructing player routes, verifying event onsets or other additional analyses (Figure 9).

**Figure 9:**
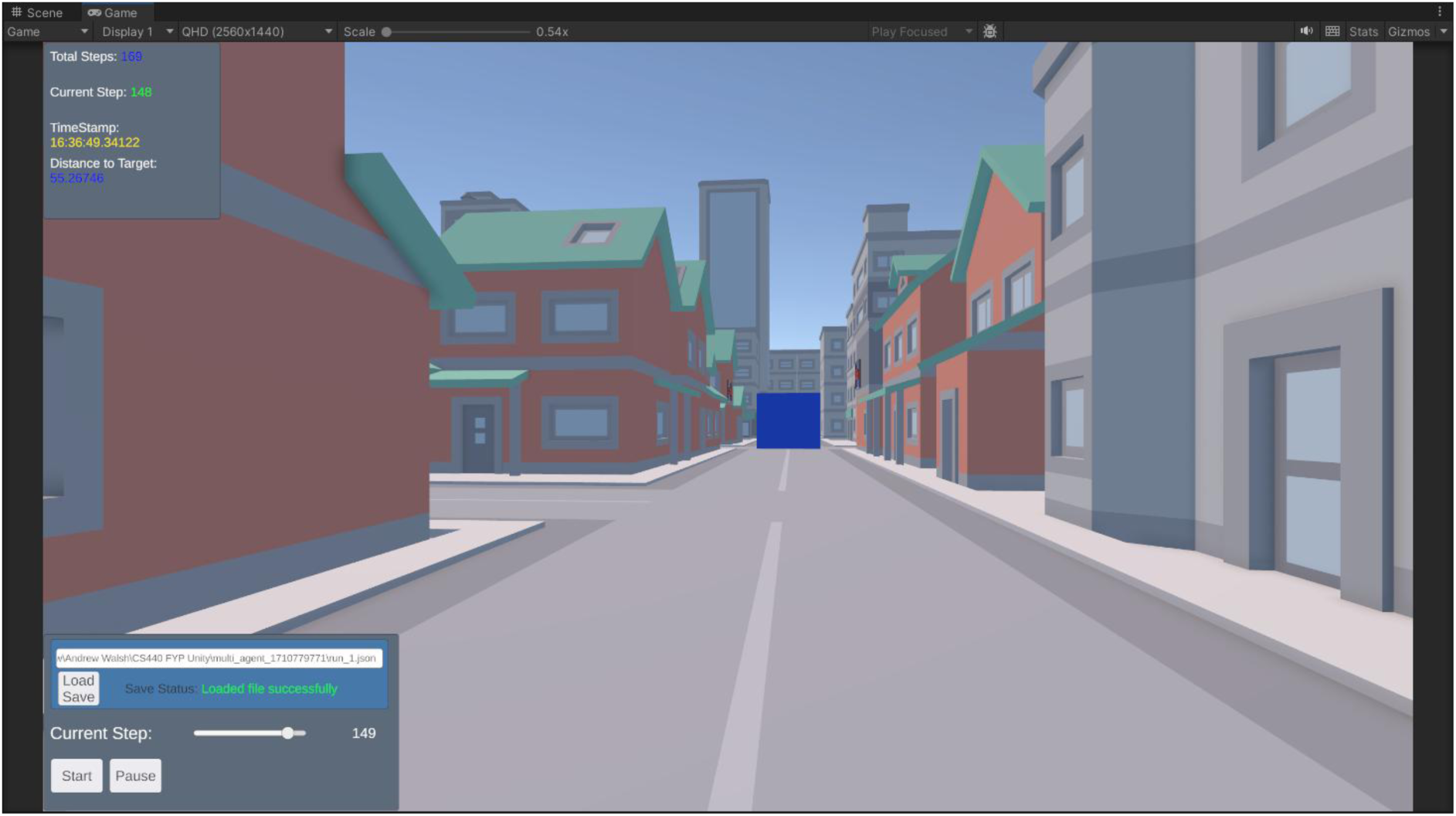
Example of a participant replay using a standard navigation task save file (.JSON). The experimenter can choose a particular step they are interested in by selecting it via the scroll bar, as well as start from the beginning or pause to observe the participant perspective during a particular step of interest.

### Adding additional tasks to WFT

1. Additional navigation trials can be added to an existing level, or a new level can be created, using the four-step procedure shown in Figure 10. Within the Unity project, the relevant task files are located at: Assets/Project/GameSystems/WayFindingTest/Tasks. *Create the new scriptable object:* In the task folder, right click -> Create -> ScriptableObjects -> WayFindingTask. This will create the task (a new trial), rename the trial.
2. *Setting up the task:* Pick the start spawn point. Then add a task by clicking the plus to add a new item to the task list. Select the **target**. Set the display text (shown at the beginning of this task in the grey instructions box). Set the task target settings (click the circle icon on the right, there should be 2 options to pick from). These settings decide how long the display text will stay on screen, how long the player has to reach the destination, and how long to wait before showing the next task. You can modify the time values of task target or create a new task target scriptable object (same method as described in Step 1 except choose WayFindingTaskTarget instead).
3. *Assign the task to the manager:* In the WayFindingTest scene, select the Manager gameobject and under the Way Finding Manager component there is a list called *Wf Task List*, click the plus icon and assign the new task created in the previous steps (it should now appear as an option to pick after clicking the circle). By default, there should be a Level 1 and Level 2.
4. *Assign the task to the UI:* To add the new task option into the UI selection at the beginning of the game, we need to add a new option to the drop down. The dropdown gameobject can be found under the following: Select the Canvas gameobject in the WayFindingTest scene, expand it to show all the child objects under the Canvas. Keep expanding until the *Task Dropdown* object can be selected. Canvas -> Main UI -> Management UI -> Panel -> Panel -> Task Dropdown. On this gameobject there is a dropdown component, at the bottom of this component there is a list of options. By default, there should be a Level 1 and Level 2 option. Click the plus icon to add another option, give it a name (ideally match it to the name of your new task). This new option will now show up in the drop down at the setup menu for the experimenter. Make sure the dropdown option is in the same order as the task in the manager component. Level 1 is the first in both lists, Level 2 is the second in both, etc. **Make sure there are no empty slots in either list.**

### Additional Technical Information, AI & Open Science

A full document containing all of the above information, as well as the technical specifications for those who are interested in designing additional or more complex tasks in the Virtual Town is available in the Supplementary Materials. However, if you wish to pick up the task and start testing, the main game file executable can be downloaded and launched on any Windows device very simply through the Zenodo link: [Link Following Acceptance]. Whilst it is possible to use Unity and follow the provided instructions in order to code our task differently, recent advances in Artificial Intelligence (AI) and Large Language Models now provide an alternative approach. We encourage using the tool as a platform, to generate additional, neuroimaging compatible versions of environments like NAVAN, that suit the researchers design, size and environmental needs. This can be done without coding efficiency, and we recommend using the code/neuroimaging compatibility from our task, to inform generative AI systems of how best to design new tasks; a developing approach in current neuroscience and psychological research (Kellmeyer, 2019). Nevertheless, we would advise that these software remain openly available (using open-source engines such as GODOT: see (Holfeld, 2023)), are not used commercially and are experimentally validated before large-scale use (Topol, 2019).

**Figure 10:**
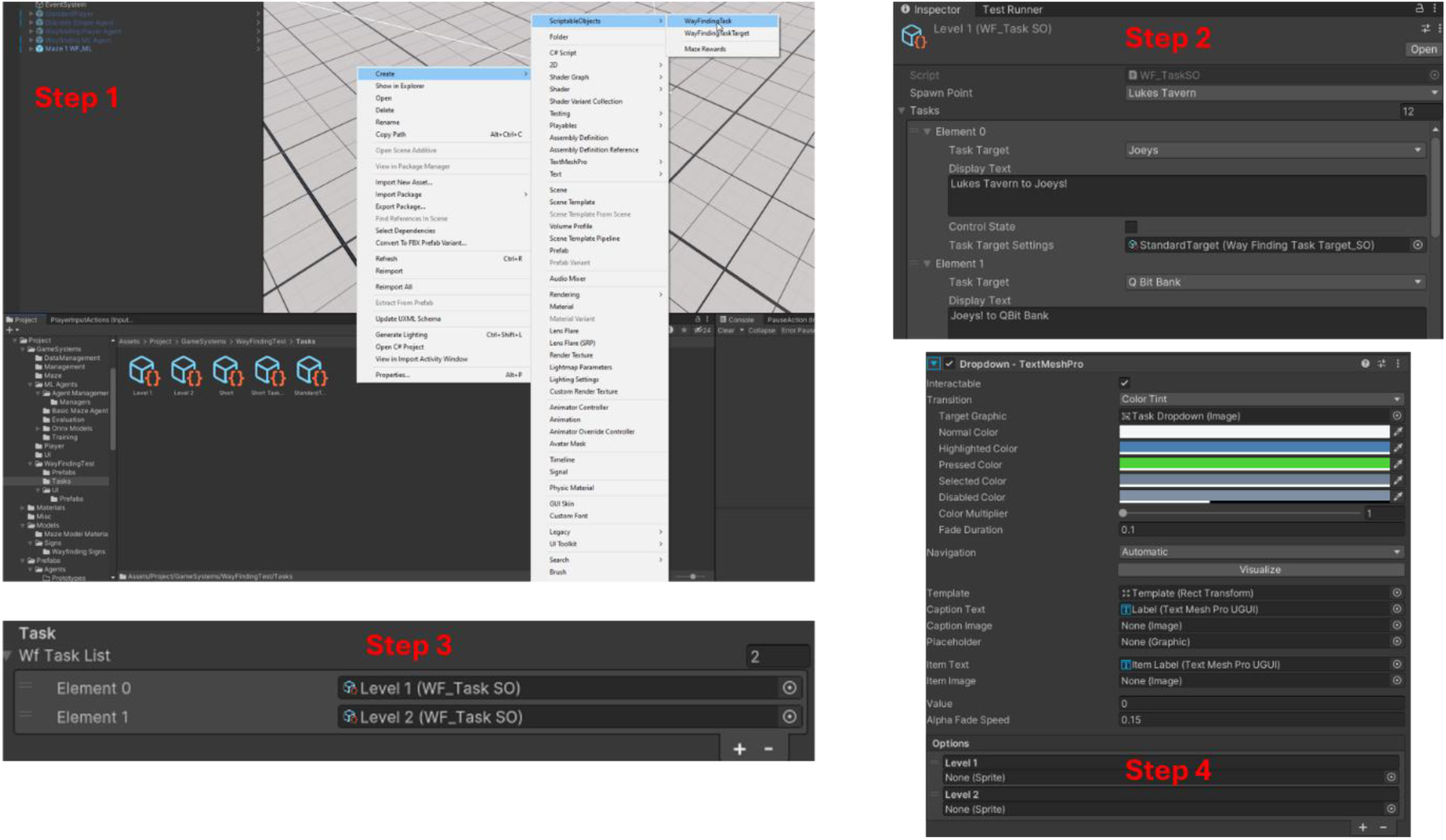
Unity workflow for creating or modifying a Wayfinding/Virtual Town task. Step 1 shows the list of scriptable objects and how the levels can be located for editing. Step 2 shows the level creation screen which can be used to edit tasks and targets within existing levels or create a new task list in a new scriptable object level. Step 3. Shows a screenshot of the task list in which new levels will be added, or altered levels will appear. Step 4 shows the screen that allows the addition of new scriptable objects and tasks to the main UI of the game as well as the order that the levels must appear in order to be successfully displayed on the main menu.

## Discussion

The current study evaluated a virtual town task (NAVAN) as a combined behavioural and neural tool for studying spatial navigation, testing three hypotheses. First, self-reported navigation skills (measured using the SBSOD) predicted navigation performance during initial exposure to the environment. SBSOD were associated with greater path efficiency, over and above age and gender, in Level 1 but not for Level 2. Second, performance improved following repeated exposure: path efficiency increased, whereas path length, standstill count, and trial duration decreased from Level 1 to Level 2. Third, as a proof-of-concept, we demonstrated that EEG activity could be recorded and synchronised with in-game events. We observed greater theta power during active navigation compared to during planning, though the small EEG sample precludes confirmatory inference. Together these findings provide preliminary evidence that this task is feasible, sensitive to individual differences and compatible with neuroimaging

### Self-assessment of Navigation Skills Reflects Initial Spatial Learning

Our first aim was to examine whether task performance related meaningfully to self-reported spatial ability as measured by the SBSOD. Several significant associations emerged, providing preliminary evidence for concurrent validity, demonstrating task feasibility and supporting our first hypothesis. SBSOD scores were negatively correlated with trial duration and path length across levels, and the strongest association was observed for path efficiency in Level 1. This relationship was not significant in Level 2, possibly due to self-assessed navigation ability being associated with initial spatial knowledge acquisition, rather than relearning or spatial knowledge updating (Lopez & Bosco, 2026). Our regression further demonstrated that SBSOD scores predicted path efficiency during Level 1 above and beyond gender and age. However, the SBSOD is a self-report measure, and associations between self-reported and objectively measured navigation performance are generally modest (Lader et al., 2025). Nevertheless, this may reflect the fact that the skills assessed by the SBSOD (e.g., orientation, mental mapping) are skills that are required for early successful encoding, but not for the relearning process that may occur in Level 2. Therefore, our task provides similar evidence to what the literature reports, but we suggest that self-assessment of spatial ability may reflect metacognition of skills required for *early* spatial learning. This could be assessed cognitively and neurophysiologically using our task by adding confidence ratings after each trial for example (Norman et al., 2019; O’Connell & Murphy, 2018).

### Evidence of Successful Spatial Navigation

Participants performed better in Level 2 than Level 1, with higher path efficiency accompanied by shorter paths, shorter trial durations, and fewer standstill periods These results support our second hypothesis and suggest that the task can be used to study the accumulation of spatial knowledge across repeated environmental exposure. The town requires participants to plan and freely navigate novel routes between landmark pairs. Therefore, this task could be used to investigate detailed mechanisms of spatial cognition and navigation in humans (Foo et al., 2005). Unlike the Virtual Water Maze, which tends to produce rapid place learning of a fixed location (Thornberry et al., 2021, 2026), we instead see gradual accumulation of environmental knowledge. This is also unlike the DSP, as there are multiple route choices participants can make in order to navigate successfully as opposed to a dual choice (Newcombe et al., 2023). It is possible that the high performance at Trial 1 of Level 1 may reflect knowledge acquired during the practice phase, which was then followed by goal combinations that were not explored during practice. However, task difficulty is matched between the two levels which is important for controlling for cognitive load and other factors that influence navigation ability (Nori et al., 2023). Therefore, whilst we found that efficiency was higher overall in Level 2, our interaction effect found that efficiency declined across trials within each level, more so in Level 1 compared to Level 2. This could be explained by spatial re-learning (Peterson et al., 2022). Participants used their existing representation of the town from Level 1 to solve the task during Level 2. However, it is possible that this could also be explained by fatigue. Considerable interindividual variability was also evident across trials (Commins et al., 2024). We suggest that this task can assist researchers with a detailed examination of individual differences in spatial navigation.

Trial order was not randomised across participants. Whilst this was a deliberate design choice to ensure a comparable learning trajectory, studies may wish to employ pseudo-randomised sequences to better isolate learning. Furthermore, assessing performance in a second exposure to the same environment in Level 2 could be interpreted as spatial relearning. This involves the correction of one’s mental schema of an environment through the acquisition of additional or new topographical knowledge (Lopez & Bosco, 2026). Because Level 2 required participants to reach previously visited landmarks from novel starting points, stored routes could not be repeated, and existing knowledge instead had to be reorganised to support new trajectories (Bécu et al., 2023; Deery & Commins, 2025). The task therefore offers a means of separating relearning from the reactivation of learned routes. Consistent with this, efficiency declined less steeply across trials in Level 2 than Level 1, which would be expected if an existing representation of the town was used to support performance. We note that this interaction was small and requires replication. Additionally, it is possible to examine mechanisms of reactivation by repeating the same landmark combinations repeatedly across time, whilst recording neural data (Jensen et al., 2024; Ólafsdóttir et al., 2018). However, whilst the grid-like nature of the virtual town supports the investigation of naturalistic human navigation in environments such as this (Coutrot et al., 2022), it reduces generalisability to other such environments. Nevertheless, this task facilitates multiple types of experimental setups that will help with answering more precise and mechanistic questions about the neural underpinnings of human spatial navigation.

### Gender Differences in Performance

Gender differences in navigation performance are well documented (Nazareth et al., 2019). Although these differences were not part of our preregistered or primary hypotheses, male participants in the present sample navigated faster and had fewer standstill periods than female participants across both levels, consistent with reported gender differences in virtual navigation performance (Munion et al., 2019). Importantly, no significant gender differences in path efficiency were observed, suggesting that male and female participants did not differ in the optimality of their routes despite differences in speed and hesitation. Well-controlled virtual translations of animal navigation tasks tend to reduce or eliminate gender differences (Coutrot et al., 2018). This suggests that researchers should incorporate the use of path efficiency as a gender-neutral index of spatial learning when using virtual tasks. We noted that male participants reported significantly higher SBSOD scores, consistent with previous work (Munion et al., 2019). This difference appears to reflect an increased self-perception of ability as well as a difference in some objective measures of navigation skills (Nazareth et al., 2019). SBSOD predicted early path efficiency after controlling for gender, indicating that path efficiency is a valid behavioural index of spatial ability regardless of self-reported navigational confidence. Coutrot et al. (Coutrot et al., 2018) reported that gender differences in virtual navigation skills tend to disappear when gender inequality is controlled for. Data collection occurred in Ireland, which has a low Gender Inequality Index (GII; (Nations, n.d.)) value. It is possible that this could provide an explanation for our findings.

### Feasibility of EEG and Other Neuroimaging Tools

The EEG component provides proof of concept that task events and continuous navigation behaviour can be synchronised with high-density EEG. Theta power was greater during active navigation relative to the planning phase, with significance at a frontal-central electrode in line with our hypothesis (Chrastil et al., 2022; Gramann et al., 2010; Javadi et al., 2019; Thornberry & Commins, 2024). We also show theta power difference at posterior electrode sites. This sample was also entirely female and are not meant to be interpreted as gender difference in neural activity. Sufficiently powered studies are needed to address these questions. Future work may employ functional magnetic resonance imaging; fMRI (Javadi et al., 2019; Spiers & Maguire, 2007) or even optically-pumped magnetometer magnetoencephalography; OPM-MEG (Roberts et al., 2019) for spatially and temporally precise neural recordings during navigation. OPMs are movement tolerant, which would facilitate accurate measurement of neural oscillations and source, alongside being able to investigate the difference between sensorimotor integration and cognitive processes during a task such as this (Lin et al., 2022). The task integration with Lab Streaming Layer provides a user-friendly approach to incorporating new neuroimaging technology. The in-text instructions and accompanying manual provide sufficient information to allow researchers to adapt the task substantially, by adding more trials or levels and/or creating blocks of trials to improve signal to noise ratio for neural recordings (Delaux et al., 2021). With nine landmarks in our task, there are a possible seventy-two unique pairings (bidirectionally) which can give three blocks of twenty-four trials before a participant repeats a route.

## Conclusions

This study provides preliminary evidence to support the use of a gamified virtual town navigation task as a behavioural and neural research tool for spatial cognition. Participants demonstrated improvements in navigational efficiency, and initial learning performance showed meaningful associations with self-reported spatial ability. The EEG proof-of-concept demonstrated that in-game events and continuous behaviour can be synchronised with neural recordings. Together, these findings support further use and validation of this task in studies of spatial cognition in healthy and clinical populations, including studies combining detailed behavioural tracking with neuroimaging or neurophysiological recording.

## Author Contributions

**C.T.:** Formal Analysis, Methodology, Data Curation, Visualization, Writing – Original Draft. **S.L.:** Conceptualization, Methodology, Software, Visualization, Writing – Original Draft. **J.K.B.B.:** Investigation, Formal Analysis, Validation, Visualization, Writing – Review & Editing. **D.M.C.:** Investigation, Formal Analysis, Validation, Visualization, Writing – Review & Editing. **L.S.:** Investigation, Formal Analysis, Writing – Original Draft. **R.O.:** Investigation, Writing – Review & Editing. **A.W.:** Methodology, Software, Writing – Review & Editing. **L.Z.:** Methodology, Software, Conceptualization, Resources, Supervision, Project Administration, Writing – Review & Editing. **T.W.:** Methodology, Software, Conceptualization, Resources, Supervision, Project Administration, Writing – Review & Editing. **A.L.V.:** Methodology, Software, Conceptualization, Resources, Supervision, Project Administration, Writing – Review & Editing. **R.W.:** Conceptualization, Resources, Supervision, Project Administration, Funding Acquisition, Writing – Review & Editing. All authors have read and agreed to the published version of the manuscript.

## Acknowledgements

We would like to thank Prof Seán Commins for his valuable insights during the initial designing of the wayfinding task.

## References

Astur, R. S., Ortiz, M. L., & Sutherland, R. J. (1998). A characterization of performance by men and women in a virtual morris water task: A large and reliable sex difference. Behavioural Brain Research, 93(1–2), 185–190. 10.1016/S0166-4328(98)00019-9

Bécu, M., Sheynikhovich, D., Ramanoël, S., Tatur, G., Ozier-Lafontaine, A., Authié, C. N., Sahel, J.-A., & Arleo, A. (2023). Landmark-based spatial navigation across the human lifespan. eLife, 12, e81318. 10.7554/eLife.81318

Van Beest, E. H., Terry, B., Booth, G., Harris, K. D., & Carandini, M. (2026). Brainwide representation of navigation (p. 2026.08.29.747979). bioRxiv. 10.64898/2026.08.29.747979

Boone, A. P., Maghen, B., & Hegarty, M. (2019). Instructions matter: Individual differences in navigation strategy and ability. Memory & Cognition, 47(7), 1401–1414. 10.3758/s13421-019-00941-5

Buzsáki, G., & Moser, E. I. (2013). Memory, navigation and theta rhythm in the hippocampal-entorhinal system. Nature Neuroscience, 16(2), 130–138. 10.1038/nn.3304

Chrastil, E. R., Rice, C., Goncalves, M., Moore, K. N., Wynn, S. C., Stern, C. E., & Nyhus, E. (2022). Theta oscillations support active exploration in human spatial navigation. NeuroImage, 262, 119581. 10.1016/j.neuroimage.2022.119581

Commins, S., Coutrot, A., Hornberger, M., Spiers, H. J., & De Andrade Moral, R. (2024). Examining individual learning patterns using generalised linear mixed models. Behavior Research Methods, 56(5), 4930–4945. 10.3758/s13428-023-02232-z

Commins, S., Duffin, J., Chaves, K., Leahy, D., Corcoran, K., Caffrey, M., Keenan, L., Finan, D., & Thornberry, C. (2020). NavWell: A simplified virtual-reality platform for spatial navigation and memory experiments. Behavior Research Methods, 52(3), 1189–1207. 10.3758/s13428-019-01310-5

Coughlan, G., Laczó, J., Hort, J., Minihane, A.-M., & Hornberger, M. (2018). Spatial navigation deficits—Overlooked cognitive marker for preclinical Alzheimer disease? Nature Reviews Neurology, 14(8), 496–506. 10.1038/s41582-018-0031-x

Coutrot, A., Manley, E., Goodroe, S., Gahnstrom, C., Filomena, G., Yesiltepe, D., Dalton, R. C., Wiener, J. M., Hölscher, C., Hornberger, M., & Spiers, H. J. (2022). Entropy of city street networks linked to future spatial navigation ability. Nature, 604(7904), 104–110. 10.1038/s41586-022-04486-7

Coutrot, A., Silva, R., Manley, E., de Cothi, W., Sami, S., Bohbot, V. D., Wiener, J. M., Hölscher, C., Dalton, R. C., Hornberger, M., & Spiers, H. J. (2018). Global Determinants of Navigation Ability. Current Biology, 28(17), 2861–2866.e4. 10.1016/j.cub.2018.06.009

Deery, R., & Commins, S. (2025). Equidistant Landmarks Fail to Produce the Blocking Effect in Spatial Learning Using a Virtual Water Maze Task with Healthy Adults: A Role for Cognitive Mapping? Brain Sciences, 15(4). 10.3390/brainsci15040414

Delaux, A., de Saint Aubert, J.-B., Ramanoël, S., Bécu, M., Gehrke, L., Klug, M., Chavarriaga, R., Sahel, J.-A., Gramann, K., & Arleo, A. (2021). Mobile brain/body imaging of landmark-based navigation with high-density EEG. European Journal of Neuroscience, 54(12), 8256–8282. 10.1111/ejn.15190

Diersch, N., & Wolbers, T. (2019). The potential of virtual reality for spatial navigation research across the adult lifespan. Journal of Experimental Biology, 222(Suppl_1), jeb187252. 10.1242/jeb.187252

Du, Y. K., Liang, M., McAvan, A. S., Wilson, R. C., & Ekstrom, A. D. (2023). Frontal-midline theta and posterior alpha oscillations index early processing of spatial representations during active navigation. Cortex, 169, 65–80. 10.1016/j.cortex.2023.09.005

Eichenbaum, H. (2017). The role of the hippocampus in navigation is memory. Journal of Neurophysiology, 117(4), 1785–1796. 10.1152/jn.00005.2017

Ekstrom, A. D., Caplan, J. B., Ho, E., Shattuck, K., Fried, I., & Kahana, M. J. (2005). Human hippocampal theta activity during virtual navigation. Hippocampus, 15(7), 881–889. 10.1002/hipo.20109

Foo, P., Warren, W. H., Duchon, A., & Tarr, M. J. (2005). Do Humans Integrate Routes Into a Cognitive Map? Map-Versus Landmark-Based Navigation of Novel Shortcuts. *Journal of Experimental Psychology: Learning*, Memory, and Cognition, 31(2), 195–215. 10.1037/0278-7393.31.2.195

Gagnon, K. T., Thomas, B. J., Munion, A., Creem-Regehr, S. H., Cashdan, E. A., & Stefanucci, J. K. (2018). Not all those who wander are lost: Spatial exploration patterns and their relationship to gender and spatial memory. Cognition, 180, 108–117. 10.1016/j.cognition.2018.06.020

Gramann, K., Onton, J., Riccobon, D., Mueller, H. J., Bardins, S., & Makeig, S. (2010). Human Brain Dynamics Accompanying Use of Egocentric and Allocentric Reference Frames during Navigation. Journal of Cognitive Neuroscience, 22(12), 2836–2849. 10.1162/jocn.2009.21369

Hegarty, M., Montello, D. R., Richardson, A. E., Ishikawa, T., & Lovelace, K. (2006). Spatial abilities at different scales: Individual differences in aptitude-test performance and spatial-layout learning. Intelligence, 34(2), 151–176. 10.1016/j.intell.2005.09.005

Hegarty, M., Richardson, A. E., Montello, D. R., Lovelace, K., & Subbiah, I. (2002). Development of a self-report measure of environmental spatial ability. Intelligence, 30(5), 425–447. 10.1016/S0160-2896(02)00116-2

Herweg, N. A., Solomon, E. A., & Kahana, M. J. (2020). Theta Oscillations in Human Memory. Trends in Cognitive Sciences, 24(3), 208–227. 10.1016/j.tics.2019.12.006

Holfeld, J. (2023). On the relevance of the Godot Engine in the indie game development industry. arXiv Preprint arXiv:2401.01909.

Iaria, G., Palermo, L., Committeri, G., & Barton, J. J. S. (2009). Age differences in the formation and use of cognitive maps. Behavioural Brain Research, 196(2), 187–191. 10.1016/j.bbr.2008.08.040

Jacobs, J. (2014). Hippocampal theta oscillations are slower in humans than in rodents: Implications for models of spatial navigation and memory. Philosophical Transactions of the Royal Society B: Biological Sciences, 369(1635), 20130304. 10.1098/rstb.2013.0304

Javadi, A.-H., Patai, E. Z., Marin-Garcia, E., Margolis, A., Tan, H.-R. M., Kumaran, D., Nardini, M., Penny, W., Duzel, E., Dayan, P., & Spiers, H. J. (2019). Prefrontal Dynamics Associated with Efficient Detours and Shortcuts: A Combined Functional Magnetic Resonance Imaging and Magnetoencenphalography Study. Journal of Cognitive Neuroscience, 31(8), 1227–1247. 10.1162/jocn_a_01414

Jensen, K. T., Hennequin, G., & Mattar, M. G. (2024). A recurrent network model of planning explains hippocampal replay and human behavior. Nature Neuroscience, 27(7), 1340–1348. 10.1038/s41593-024-01675-7

Kellmeyer, P. (2019). Artificial Intelligence in Basic and Clinical Neuroscience: Opportunities and Ethical Challenges. Neuroforum, 25(4), 241–250. 10.1515/nf-2019-0018

Kothe, C., Shirazi, S. Y., Stenner, T., Medine, D., Boulay, C., Grivich, M. I., Artoni, F., Mullen, T., Delorme, A., & Makeig, S. (2025). The lab streaming layer for synchronized multimodal recording. Imaging Neuroscience, 3, IMAG.a.136. 10.1162/IMAG.a.136

Lader, J. L., Nguyen, K. V., & Newcombe, N. S. (2025). Individual differences in navigation skill: Towards reliable and valid measures. Cognitive Research: Principles and Implications, 10(1), 27. 10.1186/s41235-025-00642-5

Lester, A. W., Moffat, S. D., Wiener, J. M., Barnes, C. A., & Wolbers, T. (2017). The Aging Navigational System. Neuron, 95(5), 1019–1035. 10.1016/j.neuron.2017.06.037

Lin, M.-H., Liran, O., Bauer, N., & Baker, T. E. (2022). Scalp recorded theta activity is modulated by reward, direction, and speed during virtual navigation in freely moving humans. Scientific Reports, 12(1), 2041. 10.1038/s41598-022-05955-9

Lopez, A., & Bosco, A. (2026). Long-term spatial memory in aging: Does research overlook retrograde memory relative to anterograde memory? Spatial Cognition & Computation, 26(1), 1–18. 10.1080/13875868.2025.2528182

Love, J., Selker, R., Marsman, M., Jamil, T., Dropmann, D., Verhagen, J., Ly, A., Gronau, Q. F., Šmíra, M., Epskamp, S., Matzke, D., Wild, A., Knight, P., Rouder, J. N., Morey, R. D., & Wagenmakers, E.-J. (2019). JASP: Graphical Statistical Software for Common Statistical Designs. Journal of Statistical Software, 88, 1–17. 10.18637/jss.v088.i02

Marchette, S. A., Bakker, A., & Shelton, A. L. (2011). Cognitive Mappers to Creatures of Habit: Differential Engagement of Place and Response Learning Mechanisms Predicts Human Navigational Behavior. Journal of Neuroscience, 31(43), 15264–15268. 10.1523/JNEUROSCI.3634-11.2011

Moffat, S. D., Hampson, E., & Hatzipantelis, M. (1998). Navigation in a ‘Virtual’ Maze: Sex Differences and Correlation With Psychometric Measures of Spatial Ability in Humans. Evolution and Human Behavior, 19(2), 73–87. 10.1016/S1090-5138(97)00104-9

Munion, A. K., Stefanucci, J. K., Rovira, E., Squire, P., & Hendricks, M. (2019). Gender differences in spatial navigation: Characterizing wayfinding behaviors. Psychonomic Bulletin & Review, 26(6), 1933–1940. 10.3758/s13423-019-01659-w

Nations, U. (n.d.). Gender Inequality Index. In Human Development Reports. United Nations. Retrieved 10 April 2026, from https://hdr.undp.org/data-center/thematic-composite-indices/gender-inequality-index

Nazareth, A., Huang, X., Voyer, D., & Newcombe, N. (2019). A meta-analysis of sex differences in human navigation skills. Psychonomic Bulletin & Review, 26(5), 1503–1528. 10.3758/s13423-019-01633-6

Newcombe, N. S., Hegarty, M., & Uttal, D. (2023). Building a Cognitive Science of Human Variation: Individual Differences in Spatial Navigation. Topics in Cognitive Science, 15(1), 6–14. 10.1111/tops.12626

Nori, R., Zucchelli, M. M., Palmiero, M., & Piccardi, L. (2023). Environmental cognitive load and spatial anxiety: What matters in navigation? Journal of Environmental Psychology, 88, 102032. 10.1016/j.jenvp.2023.102032

Norman, E., Pfuhl, G., Sæle, R. G., Svartdal, F., Låg, T., & Dahl, T. I. (2019). Metacognition in Psychology. Review of General Psychology, 23(4), 403–424. 10.1177/1089268019883821

O’Connell, R. G., & Murphy, P. R. (2018). U-turns in the brain. Nature Neuroscience, 21(4), 461–462. 10.1038/s41593-018-0122-4

O’Keefe, J., & Nadel, L. (1978). The hippocampus as a cognitive map. Clarendon Press.

Ólafsdóttir, H. F., Bush, D., & Barry, C. (2018). The Role of Hippocampal Replay in Memory and Planning. Current Biology, 28(1), R37–R50. 10.1016/j.cub.2017.10.073

Padilla, L. M., Creem-Regehr, S. H., Stefanucci, J. K., & Cashdan, E. A. (2017). Sex differences in virtual navigation influenced by scale and navigation experience. Psychonomic Bulletin and Review, 24(2), 582–590. 10.3758/s13423-016-1118-2

Peterson, M., Mead, P., Kelly, S., Esser-Adomako, E., & Blumberg, E. (2022). Spatial context target relearning following a target relocation event: Not mission impossible. *Attention, Perception*, & Psychophysics, 84(2), 474–489. 10.3758/s13414-022-02443-0

Renard, Y., Lotte, F., Gibert, G., Congedo, M., Maby, E., Delannoy, V., Bertrand, O., & Lécuyer, A. (2010). OpenViBE: An Open-Source Software Platform to Design, Test, and Use Brain–Computer Interfaces in Real and Virtual Environments. Presence, 19(1), 35–53. 10.1162/pres.19.1.35

Roberts, G., Holmes, N., Alexander, N., Boto, E., Leggett, J., Hill, R. M., Shah, V., Rea, M., Vaughan, R., Maguire, E. A., Kessler, K., Beebe, S., Fromhold, M., Barnes, G. R., Bowtell, R., & Brookes, M. J. (2019). Towards OPM-MEG in a virtual reality environment. NeuroImage, 199, 408–417. 10.1016/j.neuroimage.2019.06.010

Rolls, E. T. (2025). Hippocampal Discoveries: Spatial View Cells, Connectivity, and Computations for Memory and Navigation, in Primates Including Humans. Hippocampus, 35(1), e23666. 10.1002/hipo.23666

Schiller, D., Eichenbaum, H., Buffalo, E. A., Davachi, L., Foster, D. J., Leutgeb, S., & Ranganath, C. (2015). Memory and Space: Towards an Understanding of the Cognitive Map. The Journal of Neuroscience, 35(41), 13904–13911. 10.1523/JNEUROSCI.2618-15.2015

Spiers, H. J., & Barry, C. (2015). Neural systems supporting navigation. *Current Opinion in Behavioral Sciences*, Cognitive Control, 1, 47–55. 10.1016/j.cobeha.2014.08.005

Spiers, H. J., & Maguire, E. A. (2007). A navigational guidance system in the human brain. Hippocampus, 17(8), 618–626. 10.1002/hipo.20298

Stoianov, I. P., Pennartz, C. M. A., Lansink, C. S., & Pezzulo, G. (2018). Model-based spatial navigation in the hippocampus-ventral striatum circuit: A computational analysis. PLOS Computational Biology, 14(9), e1006316. 10.1371/journal.pcbi.1006316

Tadel, F., Baillet, S., Mosher, J. C., Pantazis, D., & Leahy, R. M. (2011). Brainstorm: A User-Friendly Application for MEG/EEG Analysis. Computational Intelligence and Neuroscience, 2011(1), 879716. 10.1155/2011/879716

Taillade, M., N’Kaoua, B., & Sauzéon, H. (2016). Age-Related Differences and Cognitive Correlates of Self-Reported and Direct Navigation Performance: The Effect of Real and Virtual Test Conditions Manipulation. Frontiers in Psychology, 6. 10.3389/fpsyg.2015.02034

Thornberry, C., Cimadevilla, J. M., & Commins, S. (2021). Virtual Morris water maze: Opportunities and challenges. Reviews in the Neurosciences, 32(8), 887–903. 10.1515/revneuro-2020-0149

Thornberry, C., Cimadevilla, J. M., & Commins, S. (2026). Virtual Morris Water Task: Procedures and Protocols for the Assessment of Spatial Navigation and Memory. Current Protocols, 6(3), e70340. 10.1002/cpz1.70340

Thornberry, C., & Commins, S. (2024). Frontal delta and theta power reflect strategy changes during human spatial memory retrieval in a virtual water maze task: An exploratory analysis. Frontiers in Cognition, 3. 10.3389/fcogn.2024.1393202

Tolman, E. C. (1948). Cognitive maps in rats and men. Psychological Review, 55(4), 189–208. 10.1037/h0061626

Topol, E. J. (2019). High-performance medicine: The convergence of human and artificial intelligence. Nature Medicine, 25(1), 44–56. 10.1038/s41591-018-0300-7

Ventura, M., Shute, V., Wright, T. J., & Zhao, W. (2013). An investigation of the validity of the virtual spatial navigation assessment. Frontiers in Psychology, 4. 10.3389/fpsyg.2013.00852

Whelan, R., Barbey, F. M., Cominetti, M. R., Gillan, C. M., & Rosická, A. M. (2022). Developments in scalable strategies for detecting early markers of cognitive decline. Translational Psychiatry, 12(1), 473. 10.1038/s41398-022-02237-w

Yavuz, E., & Spiers, H. J. (2026). Determinants of individual navigation ability. Nature Reviews Psychology, 5(4), 273–289. 10.1038/s44159-026-00544-3

