## Supplementary Materials for "An open-access neuroimaging-compatible virtual town task for assessing human spatial navigation"

### Correlation Analyses

*Supplementary Table 1: Pearson's Correlations* for all available variables and SBSOD.
Note: L1 represents Level 1 and L2 represents Level 2.

| | | | Pearson's r | p |
|---|---|---|---|---|
| SBSOD | - | trial_duration_L1 | -0.349* | .010 |
| SBSOD | - | trial_duration_L2 | -0.208 | .136 |
| SBSOD | - | path_length_L1 | -0.424** | .002 |
| SBSOD | - | path_length_L2 | -0.346* | .011 |
| SBSOD | - | path_efficiency_L1 | 0.460*** | < .001 |
| SBSOD | - | path_efficiency_L2 | 0.264 | .056 |
| SBSOD | - | standstill_count_L1 | -0.196 | .160 |
| SBSOD | - | standstill_count_L2 | -0.100 | .475 |

* p < .05, ** p < .01, *** p < .001

**Supplementary Figure 1:** Pearson correlation plots for all available variables and SBSOD scores to examine preliminary relationships and check for multicollinearity.

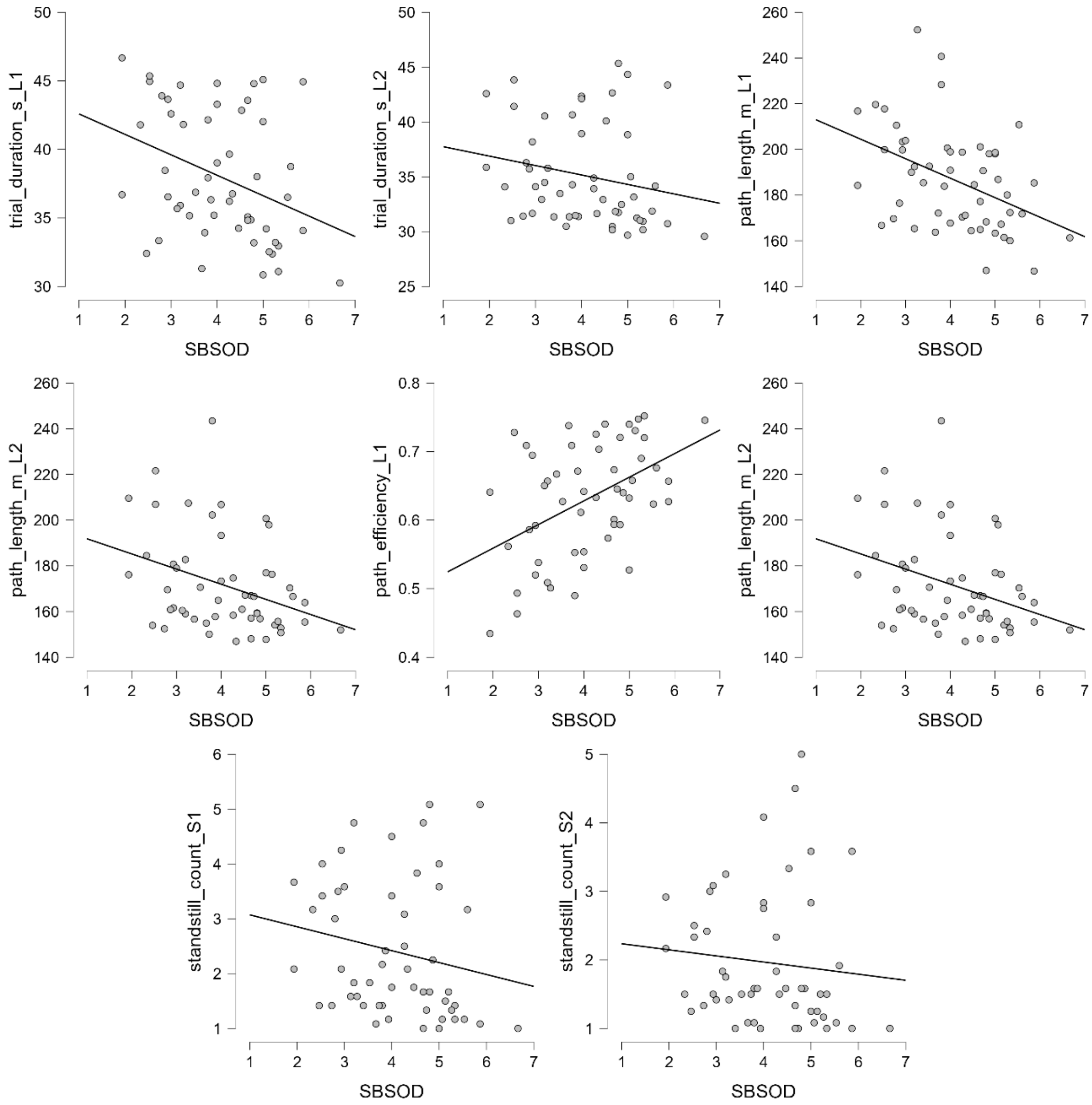
